# Integrating virtual cell modeling with patient-derived transcriptomics to uncover target signals in rheumatoid arthritis

**DOI:** 10.64898/2026.04.19.719532

**Authors:** Kazuki Nakanishi, Hideyuki Shimizu

## Abstract

Rheumatoid arthritis affects millions of people worldwide, yet a substantial fraction of patients fail to achieve lasting benefit from current therapies. AI-based virtual cell models offer a promising route to simulate the effects of thousands of potential drug targets computationally, but whether their predictions genuinely reflect what happens in the disease-relevant cells of patients undergoing treatment has been unclear. Here, we introduce ImmunoSTATE, a clinically anchored framework that benchmarks empirical and virtual perturbations directly against patient-derived treatment trajectories in pathogenic CXCL13+ synovial T cells from nine patients with rheumatoid arthritis sampled before and after treatment. Empirical genome-wide CRISPR interference screening prioritizes T cell receptor (TCR)-proximal signaling over the measurable components of the JAK-STAT pathway for transcriptomic reversal of the disease signature, a result that remained robust across leave-one-patient-out reanalyses. Comparison of virtual perturbation signals across healthy donor CD4+ T cells and rheumatoid-arthritis patient cells reveals broad preservation of target priorities, together with selected disease-context-enriched shifts, including stronger prioritization of SOX4 in patient cells. ImmunoSTATE provides a clinically anchored framework for evaluating virtual cell models in disease-relevant immune contexts and demonstrates that distinguishing training-set status and cellular context are essential steps in the reliable interpretation of virtual perturbation outputs.

## Introduction

Rheumatoid arthritis (RA) is a chronic autoimmune disorder characterized by dysregulated immune circuits that sustain persistent synovial inflammation and tissue destruction^1^. Although Janus kinase (JAK) inhibitors have revolutionized the therapeutic landscape by targeting intracellular signaling cascades downstream of multiple pro-inflammatory cytokines^2,3^, a substantial fraction of patients fail to achieve durable clinical benefit. This gap raises a fundamental question of whether molecular targets beyond those already exploited by existing drugs could more effectively reverse disease-associated cell states in RA. Traditional target identification has relied on candidate-driven approaches focused on canonical signaling pathways^1^. However, the complexity of synovial immune cell states, particularly the emergence of pathogenic peripheral helper-like T cell populations^4,5,6^, demands a more systematic and unbiased evaluative framework.

Recent experimental advances have made this question increasingly approachable by bridging the gap between high-throughput perturbation screening and clinical observation. Genome-scale CRISPR interference (CRISPRi) atlases in primary human CD4+ T cells^7^ now provide comprehensive maps of how silencing each of thousands of genes affects cell state. Simultaneously, single-cell RNA sequencing (scRNA-seq) of clinical specimens enables direct measurement of treatment-associated shifts in gene expression within disease-relevant compartments such as the inflamed synovium or synovial fluid^8,9^. Together, these resources allow the systematic comparison of experimentally observed perturbation effects with the gene expression changes that occur in patient cells before and after treatment. Such comparisons go beyond relying solely on known signaling pathways or the historical record of which drugs have been developed, and are essential for validating whether the molecular signatures of *in vitro* gene knockdown truly mirror the transcriptomic changes observed in patients undergoing successful therapy.

In parallel, AI-based computational models have been developed to predict how individual cells respond to the silencing of specific genes, offering a scalable alternative to exhaustive laboratory screening^10,11^. One such model, STATE^12^, uses a large language model-style architecture trained on millions of single-cell perturbation measurements to simulate gene-silencing effects directly within a given cellular context. By learning patterns from large experimental datasets, these models can in principle predict the effects of many perturbations that would be difficult to test experimentally. However, recent studies have shown that these models often struggle to generalize accurately, particularly for genes not well represented in their training data^10, 11^. Critically, direct evaluation against patient-derived therapeutic transcriptomic shifts remains limited, and it is often unclear whether signals observed in healthy donor perturbation datasets are preserved in disease-relevant patient cells.^13^

Here, we introduce ImmunoSTATE, a framework that integrates patient-derived scRNA-seq, genome-scale CRISPRi screening, and virtual perturbation modeling to benchmark therapeutic target prioritization in the pathogenic T cells found in the joints of RA patients. Using gene expression profiles from nine patients sampled before and after treatment as a clinical reference, we first asked which experimental perturbations most closely recapitulate the transcriptomic direction of clinical improvement. We then asked whether virtual perturbation modeling recovered interpretable target-specific signal directly in pre-treatment patient cells, and whether those virtual signals were preserved across healthy and disease contexts, and explicitly considered which outputs should and should not be interpreted as biologically comparable. Empirical screening consistently identified T cell receptor (TCR)-proximal signaling, the molecular machinery that activates T cells in response to antigen recognition, as having greater potential to reverse the RA disease signature than the JAK-STAT pathway currently targeted by standard-of-care drugs^2,3^, with signaling molecules such as *ZAP70*^14^ and *CD3E*^15^ ranking among the strongest candidates genome-wide. In the virtual arm, multiple trained targets aligned positively with the patient-derived treatment direction, while comparison across healthy donor and rheumatoid-arthritis cells revealed both shared signal and selected rheumatoid-arthritis-context-enriched shifts. Together, these findings establish ImmunoSTATE as a clinically anchored approach for evaluating virtual cell models and for distinguishing shared from disease-context-enriched target signals in immune-mediated disease.

## Results

### A clinically anchored framework for benchmarking therapeutic perturbations in rheumatoid arthritis

To establish a rigorous benchmark for therapeutic target prioritization, we integrated longitudinal patient-derived transcriptomics with multi-modal perturbation screening (**Fig. 1**). Conventional drug discovery efforts often rely on healthy donor cells or immortalized cell lines, which inadequately represent the dysregulated molecular landscape of the inflamed autoimmune joint^1^. We therefore analyzed single-cell RNA-sequencing (scRNA-seq) profiles from the synovial fluid of nine patients with rheumatoid arthritis (RA) undergoing treatment with either the Janus kinase (JAK) inhibitor tofacitinib^2^ (*n* = 4) or the tumor necrosis factor (TNF) inhibitor adalimumab^3^ (*n* = 5). Our analysis focused on CXCL13+ synovial T cells^16^, a highly pathogenic population implicated in B cell help and synovial inflammation. This procedure yielded 1,307 pre-treatment and 3,600 post-treatment cells. Consistent with a pathogenic helper-like and dysfunctional phenotype, pre-treatment CXCL13+ T cells expressed *LAG3*^17^, *CTLA4*^18^, *HAVCR2*^19^, and *ENTPD1*^20^ at higher levels relative to the remaining pre-treatment CD4+ T cells (**Fig. 1A**). These cells were used to define the clinical benchmark (**Fig. 1B)**.

**Figure 1.**
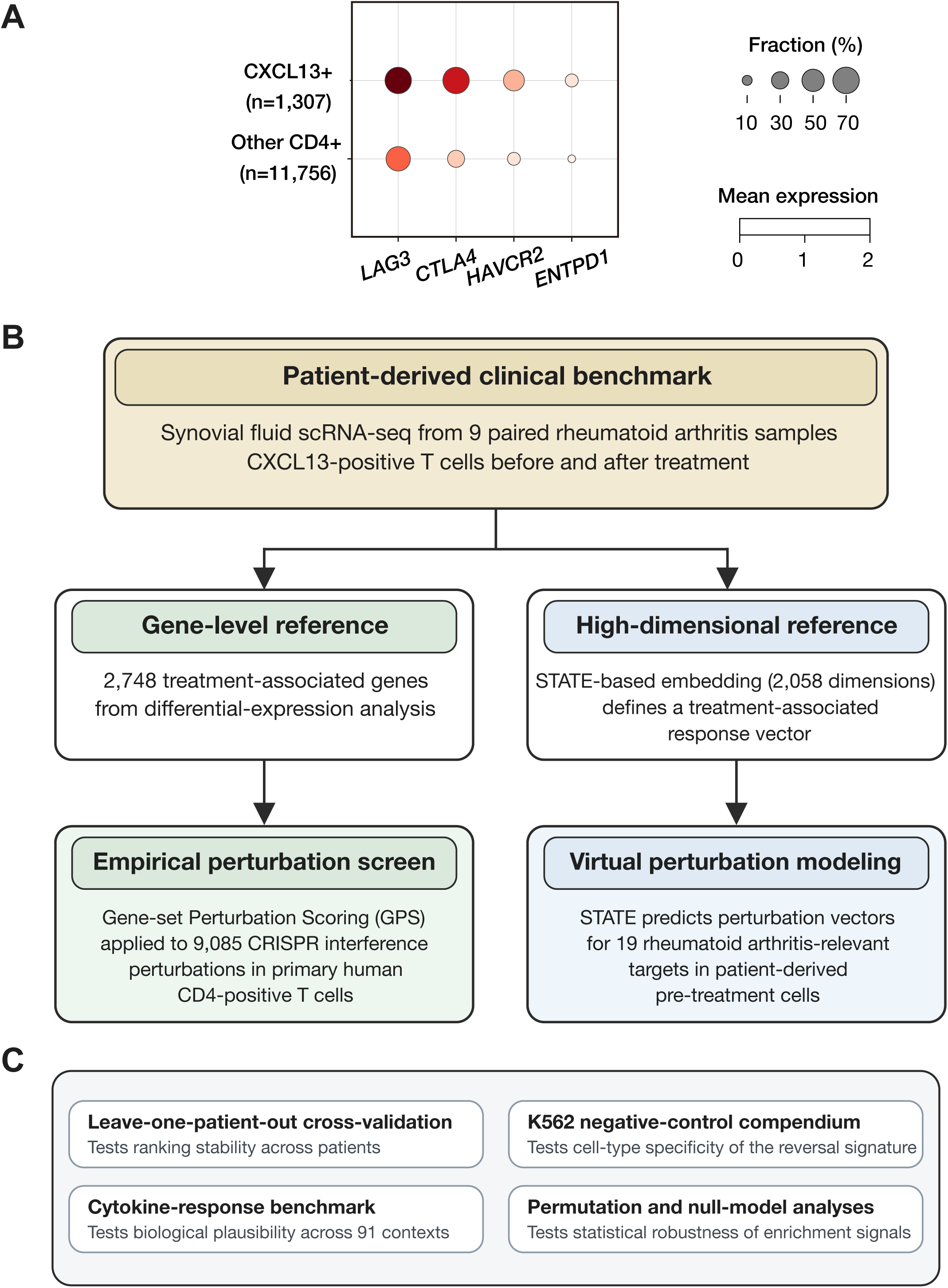
Clinically anchored study design and robustness controls. **A,** CXCL13⁺ T cells express canonical exhaustion and T peripheral helper markers. Dot plot showing expression of *LAG3, CTLA4, HAVCR2*, and *ENTPD1* in pre-treatment CXCL13⁺ pathogenic T cells (n = 1,307) versus all other pre-treatment CD4⁺ T cells (n = 11,756) pooled across the nine patients. Dot color indicates mean log-normalized expression per population and dot size indicates the fraction of cells with non-zero expression. CXCL13⁺ cells show higher expression and a greater expressing fraction for all four markers, consistent with the established peripheral helper-like and dysfunctional T cell phenotype. **B,** Study workflow. Single-cell RNA-sequencing (scRNA-seq) data from synovial fluid CXCL13^+^ T cells of nine rheumatoid arthritis (RA) patients sampled before and after treatment were used to define two patient-derived clinical references. The first is a 2,748-gene differential-expression signature serving as a gene-level reference for Gene-set Perturbation Scoring (GPS). The second is a treatment-associated response vector defined in the 2,058-dimensional SE-600M embedding space. The gene-level signature was compared with genome-wide CRISPR interference (CRISPRi) perturbations from a primary human CD4^+^ T cell atlas using GPS. In parallel, the response vector was compared with virtual perturbation vectors generated by the STATE model for 19 RA-relevant targets applied directly to patient-derived pre-treatment cells. **C,** Robustness and validation controls. The four boxes summarize four checks of the main result. Leave-one-patient-out analysis gave similar rankings across the nine re-analyses. A permutation test with 10,000 random shuffles remained significant after correction for multiple comparisons. Comparison with an unrelated K562 leukemia-cell screen showed only 4 overlapping genes among the top 100 hits, showing that the result was specific to T cells. In the Schmidt cytokine-conditioned perturbation benchmark, 7 of 8 clinically relevant contexts followed the expected directional pattern, supporting biological concordance of the empirical prioritization framework.

One component of this benchmark was a gene-level reference consisting of 2,748 genes that changed significantly after therapy by differential-expression analysis^21^. The second was a high-dimensional reference defined in SE-600M embedding space, the 2,058-dimensional single-cell representation used within the STATE framework^12^. In this space, we calculated the mean latent-space shift from pre-treatment to post-treatment cells across the clinical cohort, yielding a patient-derived treatment-associated response vector that captured the shared molecular trajectory of successful therapy irrespective of drug class. Throughout this study, we define transcriptomic reversal as a perturbation that shifts pre-treatment cells toward the post-treatment state.

Using these two clinical references, we implemented a dual-screening strategy to evaluate candidate therapeutic targets (**Fig. 1B**). First, Gene-set Perturbation Scoring (GPS) was utilized to quantify the empirical transcriptomic reversal potential of 9,085 genome-wide CRISPRi perturbations from a primary human CD4+ T cell atlas^7^, scoring each knockdown with the Reversal Gene Expression Score (RGES)^22^, which measures how strongly a given perturbation shifts the transcriptome toward the post-treatment reference. Second, the STATE virtual cell model^12^ was used to generate *in silico* perturbation vectors for 19 curated RA-relevant targets applied directly to the patient-derived pre-treatment cells (**Supplementary Table S1**). Thus, the empirical screen assessed experimentally measured perturbation effects across the genome, whereas the virtual screen evaluated model-predicted effects for a focused set of disease-relevant targets in the patient-cell context.

To test the robustness and biological specificity of this framework, we next addressed four questions summarized in **Figure 1C**: whether the results remained stable after removal of any individual patient, whether the observed signal could arise by chance, whether the same scoring procedure would produce a similar signal in an unrelated cell type, and whether target ranking agreed with an independent cytokine-conditioned perturbation benchmark. These analyses supported a stable, T cell-specific, and biologically plausible signal. The unrelated-cell-type comparison used K562 cells^23^, a chronic myeloid leukemia cell line distinct from synovial T cells, whereas the independent benchmark was based on 91 cytokine stimulation conditions in primary human T cells^24^. In addition, nine-fold leave-one-patient-out cross-validation confirmed that the population-level findings were not driven disproportionately by any single patient.

### Empirical screening prioritizes TCR-proximal signaling as a superior axis for transcriptomic reversal

To identify the most potent molecular targets for normalizing the pathogenic transcriptional state of RA synovial T cells, we conducted an unbiased genome-wide evaluation of transcriptomic reversal potential. Although Janus kinase (JAK) inhibitors are a cornerstone of current RA therapy^2,3^, it remains unclear whether they represent the theoretical optimum for reversing the disease-specific transcriptomic signature of CXCL13+ T cells. We addressed this by applying GPS to a primary human CD4+ T cell CRISPRi atlas encompassing 9,085 robust perturbations^7^, ranking each gene knockdown by its RGES^22^, a metric that quantifies the degree to which silencing a given gene shifts the cellular transcriptome toward the post-treatment clinical reference.

The genome-wide GPS ranking consistently identified TCR-proximal signaling^25^ as the strongest pathway-level axis for reversal of the RA transcriptomic program (**Fig. 2A, Supplementary Fig. S1**). The top-ranked genes included multiple canonical TCR-proximal components, including ZAP70^14^, CD3E^15^, CD3G^15^, and LCP2^26^, together with the mitochondrial/bioenergetic hit COX11^27^, and seven of the nine evaluated TCR-proximal genes ranked within the top 50 perturbations genome-wide. This pattern highlighted a dominant TCR-proximal cluster among the strongest reversal-associated perturbations. At the pathway level, TCR-proximal genes achieved a median GPS rank of 6 out of 9,085, placing the entire pathway among the strongest reversal candidates genome-wide. This contrasted sharply with the 18 measurable JAK-STAT pathway genes, which achieved a median rank of 8,166 (**Fig. 2B**, *p* = 6.4 × 10⁻⁵). Among measurable JAK-STAT components, only *STAT3* entered the top 100 (rank 25), whereas the remaining 17 measurable members all ranked 1,457th or lower, including clinically familiar genes such as *JAK2* and *TYK2*, which ranked 4,623rd and 8,636th, respectively.

**Figure 2.**
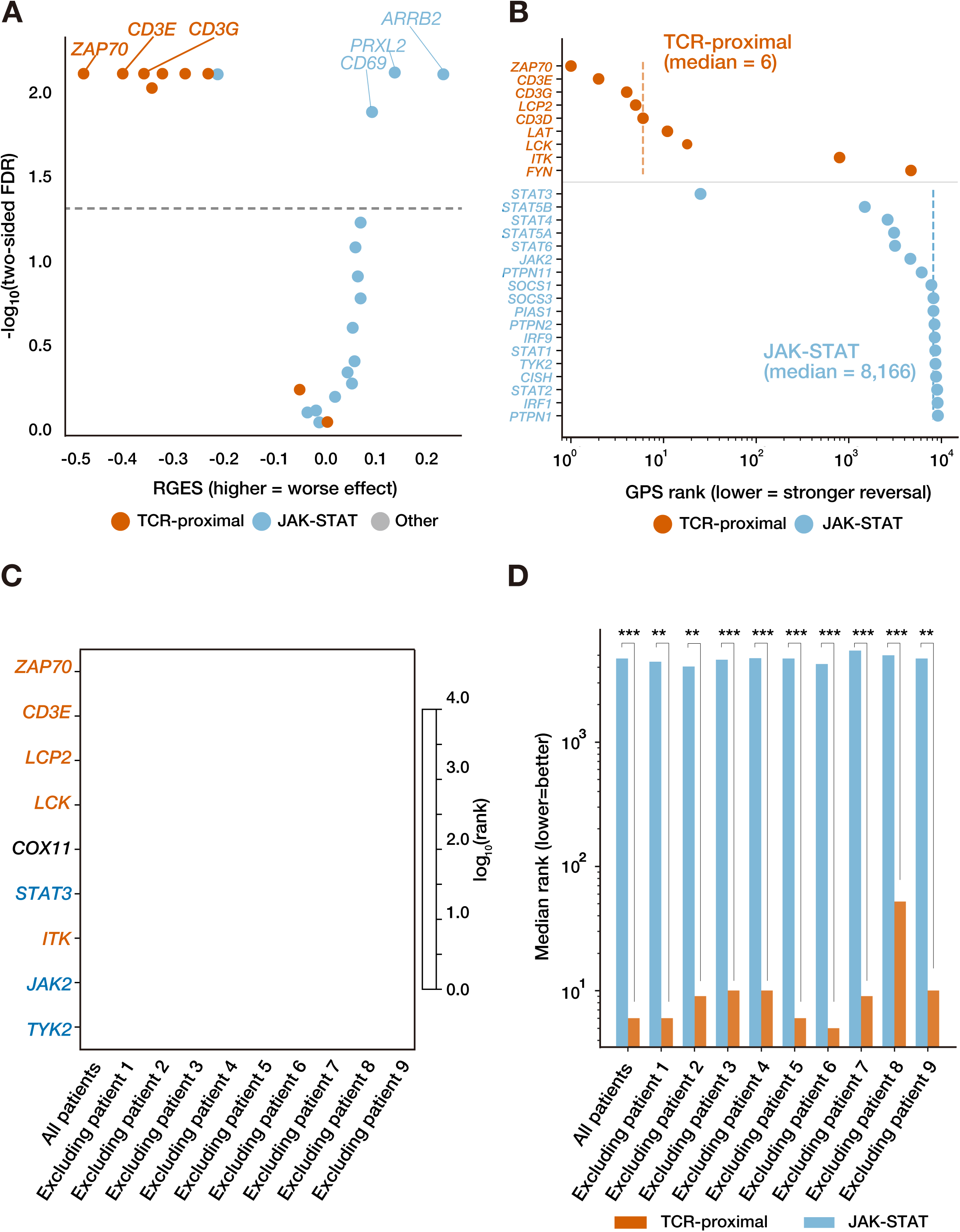
Empirical screening prioritizes TCR-proximal signaling over the measurable JAK-STAT pathway. **A,** Two-sided descriptive overview of the genome-wide perturbation screen. Each point represents one of 9,085 CRISPRi perturbations in primary human CD4+ T cells. The x-axis shows the Reversal Gene Expression Score (RGES), where negative values indicate movement toward the post-treatment RA reference and positive values indicate movement away from it. The y-axis shows a two-sided false-discovery rate recalculated from absolute effect magnitude so that perturbations on both sides of zero are visible. Orange points denote TCR-proximal genes, blue points denote measurable JAK-STAT genes, and gray points denote all other perturbations. The dashed horizontal line marks false-discovery rate threshold of 0.05. A reversal-oriented one-sided view of this distribution, used for therapeutic ranking, is provided in Supplementary Figure S1. **B,** Direct pathway-level comparison of GPS ranks. Genome-wide GPS ranks for canonical TCR-proximal genes (orange) and measurable JAK-STAT genes (blue) are plotted on a logarithmic scale, where lower rank indicates stronger transcriptomic reversal. Dashed vertical lines indicate the median rank for each pathway. TCR-proximal genes have achieved a median rank of 6, whereas measurable JAK-STAT genes achieved a median rank of 8,166 (Mann-Whitney U test, p = 6.4 × 10^&’^). *JAK1* and *JAK3* were absent from this analysis as they could not be measured in the primary CD4+ T cell CRISPRi atlas due to their essentiality for T cell survival. **C,** Stability of target ranking across the clinical cohort. A heatmap displays the log_10_ transformed GPS ranks for selected therapeutic targets derived from all nine patients combined and from each of the nine leave-one-patient-out (LOPO) cross-validation folds. Lighter color intensity indicates a higher ranking and stronger disease-signature reversal potential. **D,** Robustness of pathway prioritization across LOPO folds. Median genome-wide GPS ranks for TCR-proximal and JAK-STAT signaling genes are shown for all nine patients combined and each individual LOPO fold. TCR-proximal signaling consistently achieves lower median ranks than JAK-STAT signaling regardless of the excluded patient. Asterisks denote the statistical significance of the pathway contrast within each fold (\*\**P* < 0.01, \*\*\**P* < 0.001; Mann-Whitney U test).

To confirm that the TCR-proximal signal was not driven by any single patient, nine-fold leave-one-patient-out reanalyses were performed. The overall target ranking remained highly consistent across all nine reanalyses (**Fig. 2C**, Kendall’s rank correlation τ^28^ = 0.761 ± 0.117), and the pathway-level contrast between TCR-proximal and JAK-STAT signaling remained significant in every fold (**Fig. 2D**, *p* < 0.002).

To confirm that the TCR-proximal signal reflected T cell-specific biology rather than a generic response to gene silencing, we applied the same RA reversal signature to an orthogonal CRISPRi dataset generated in K562 chronic myeloid leukemia cells^23^, a cell line unrelated to synovial T cells. Only 4 of the top 100 hits were shared between the K562 and primary T cell screens (**Supplementary Fig. S2A**), with the leading K562 perturbations dominated by RNA-processing factors and Integrator-complex members rather than immune signaling genes (**Supplementary Fig. S2B**).

To assess biological coherence, we benchmarked the RA reversal signature against the Schmidt cytokine-conditioned CRISPRi resource^24^, reasoning that pro-inflammatory cytokine blockade contexts should rank favorably whereas anti-inflammatory stimulation contexts should rank at the opposite end. Among eight prespecified clinically relevant contexts, seven followed this expected directional pattern (**Supplementary Fig. S3A**), and the observed concordance exceeded chance expectation under three of five null models (**Supplementary Fig. S3B**).

Together, these results establish that TCR-proximal signaling more closely recapitulates the transcriptomic direction of patient improvement than the measurable components of the JAK-STAT pathway, and that this finding is stable across patients, specific to the T cell context, and biologically plausible.

### ImmunoSTATE reveals interpretable virtual-perturbation signal in rheumatoid-arthritis patient cells

Having established the empirical landscape, we next applied virtual perturbation modeling to assess whether target-specific signal was recoverable directly in pre-treatment RA cells. Using the patient-derived treatment-associated response vector as a clinical reference, we calculated the cosine similarity between each ImmunoSTATE-predicted perturbation vector^12^ and the mean pre-treatment-to-post-treatment shift in the native 2,058-dimensional SE-600M embedding space. Positive values indicate agreement with the treatment-associated shift.

Among the 16 targets for which the STATE model had direct training data, 10 showed significant positive alignment (**Fig. 3**). Targets with significant positive alignment included *IRF4*^29^ and *SOX4*^30^ among transcriptional regulators, and *ITK*^31^ and *SYK*^32^ among TCR-proximal signaling kinases, as well as *NFKB1*^33^, *EOMES*^34^, *MAF*^35^, *TOX*^36^, *STAT3*^37^, and *BTK*^38^. By contrast, *STAT4*^39^, *TBX21*^40^, *RORC*^41^, and *TOX2*^42^ showed negative alignment. These results indicate that interpretable, target-specific virtual signals for promising therapeutic candidates are robustly recoverable by ImmunoSTATE directly in rheumatoid-arthritis patient cells.

**Figure 3.**
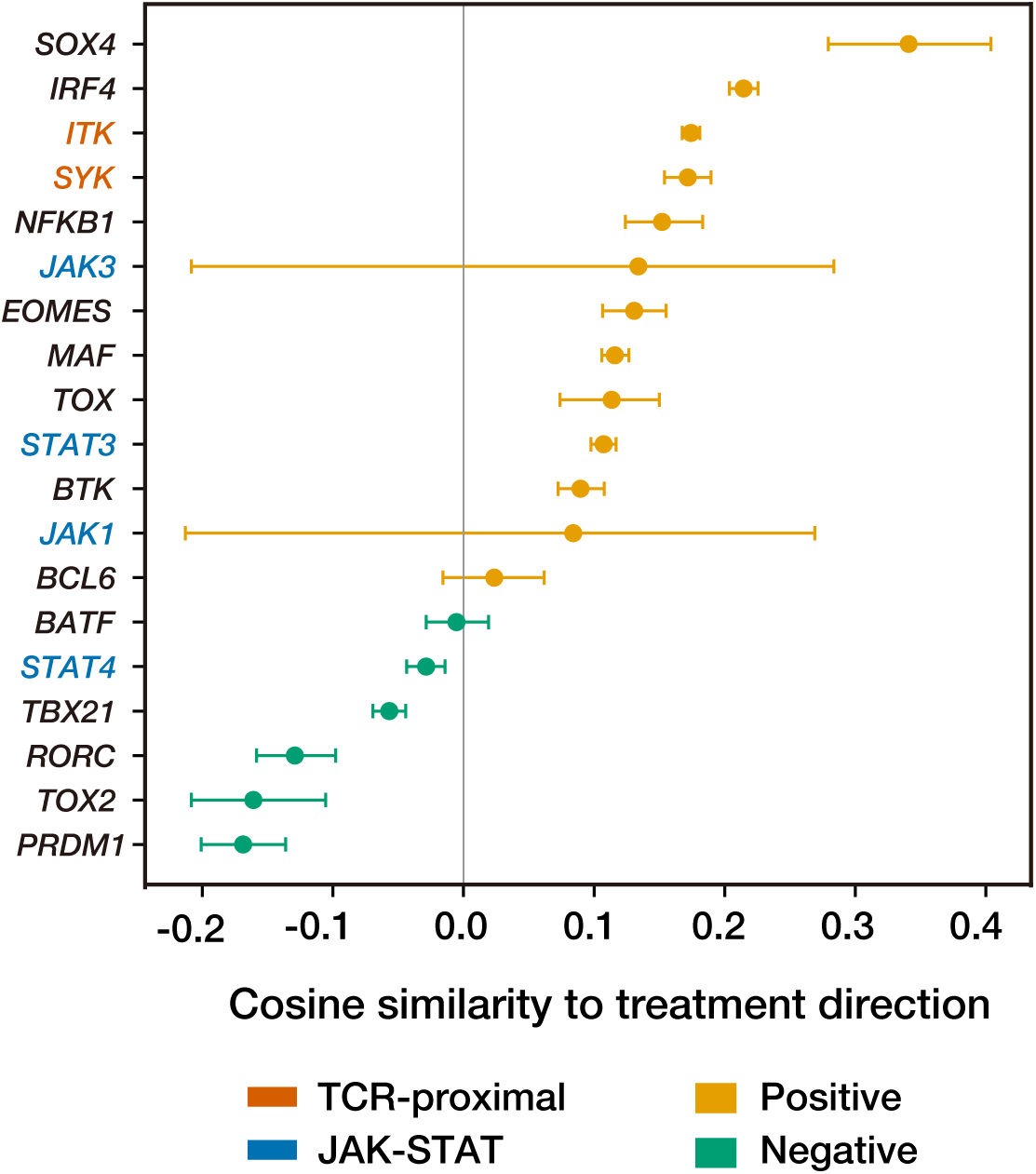
ImmunoSTATE reveals interpretable virtual-perturbation signal in rheumatoid-arthritis patient T cells. Cosine similarity between each STATE-predicted perturbation vector and the patient-derived treatment-associated response vector, computed in the native 2,058-dimensional SE-600M embedding space. Positive values indicate directional alignment with the clinical treatment shift. Points indicate mean cosine similarity estimated from 2,000 bootstrap iterations and error bars denote 95% confidence intervals. Gene name color indicates pathway membership: orange, TCR-proximal; blue, JAK-STAT. Marker color indicates estimate status: gold, positive estimate; green, negative estimate. Among the 16 targets represented in the model training distribution, 10 showed significant positive alignment, including *SOX4* and *IRF4* among transcriptional regulators and *ITK* and *SYK* among TCR-proximal signaling kinases. *JAK1, JAK3,* and *PRDM1* were absent from the model training set and are shown for transparency; these targets were not used for comparative interpretation with trained targets.

### Virtual prioritization is broadly preserved across contexts but reveals selected rheumatoid-arthritis-context shifts

We next asked whether ImmunoSTATE prioritization in rheumatoid-arthritis patient cells simply recapitulated a generic CD4+ T-cell prioritization profile or instead reflected the biology of the CXCL13+ inflammatory helper state. To isolate cellular context, we applied the same ImmunoSTATE scoring procedure to healthy donor CD4+ T cells and to patient-derived rheumatoid-arthritis CXCL13+ T cells, and compared the 16 targets represented in the model training distribution (**Fig. 4**). Target rankings were significantly concordant across contexts (Spearman ρ = 0.65, p = 0.006), indicating that a substantial component of the virtual prioritization structure reflects shared CD4+ T-cell biology. However, the rheumatoid-arthritis patient-cell profile was not a simple copy of the healthy-cell profile. *SOX4* showed the clearest rheumatoid-arthritis-context amplification, consistent with its known role in promoting CXCL13-producing helper T cells under inflammatory conditions. IRF4 and ITK remained strong in both contexts, consistent with a context-stable T-cell activation and T-cell-receptor-proximal axis. By contrast, *STAT4*, *RORC*, *TOX2*, and *BCL6*^43^ were attenuated in rheumatoid-arthritis patient cells despite stronger scores in healthy donor CD4+ T cells, suggesting that conventional Th1, Th17, or follicular-helper-like programs are down-weighted in the CXCL13+ disease-cell context. Thus, ImmunoSTATE separates shared CD4+ T-cell reversal biology from patient-cell-context reweighting, highlighting a SOX4-associated CXCL13+ helper-cell axis that would be underestimated from healthy-cell evaluation alone.

**Figure 4.**
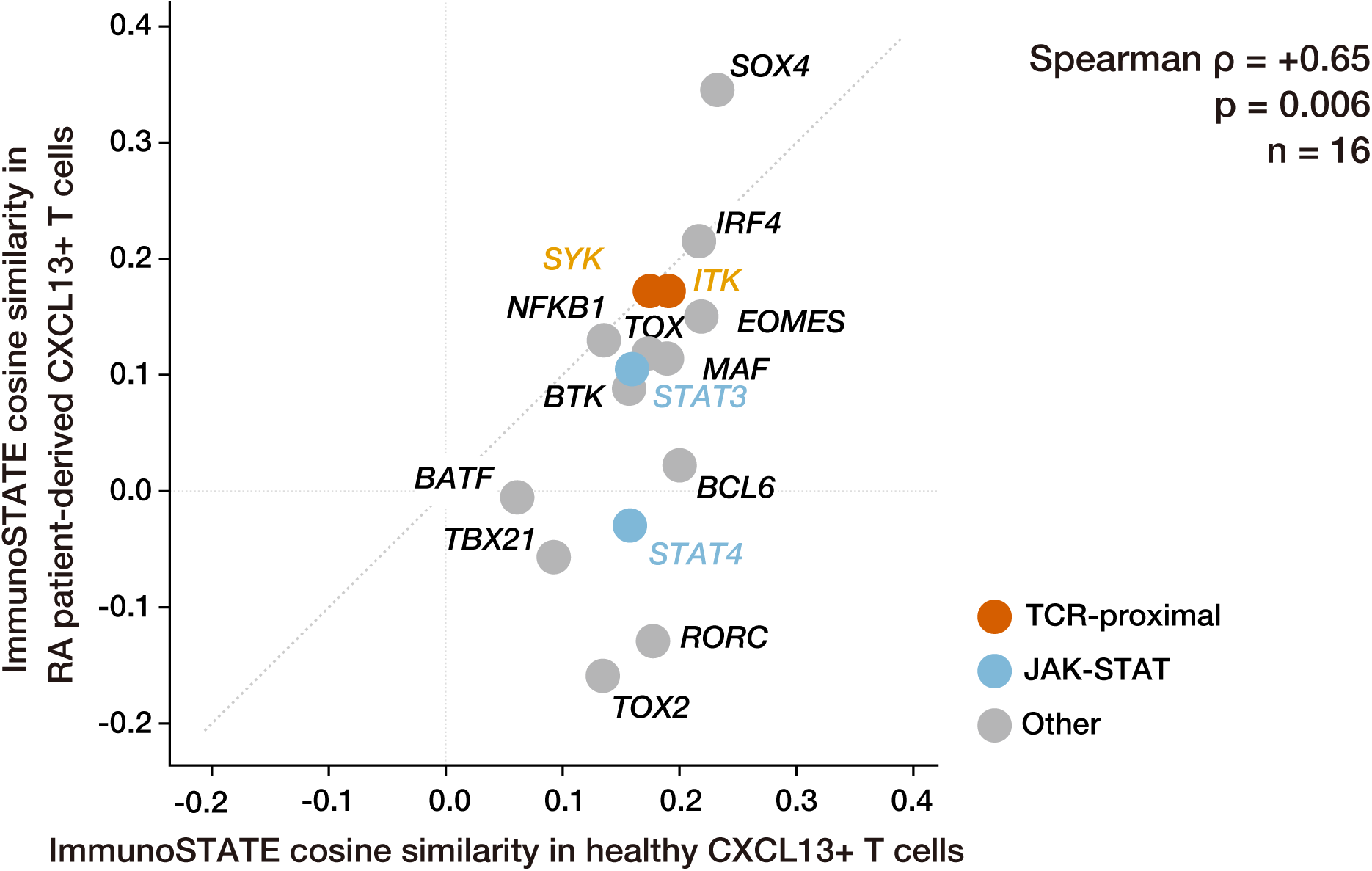
Virtual prioritization is broadly preserved across contexts but reveals selected rheumatoid-arthritis-context shifts. ImmunoSTATE virtual perturbation vectors were computed separately in healthy donor CD4+ T cells and in rheumatoid-arthritis patient CXCL13+ T cells. In both contexts, target scores were calculated as cosine similarity to the same patient-derived treatment-associated reference direction derived from GSE296117 rheumatoid-arthritis CXCL13+ T cells. In both axes, positive values indicate directional alignment with the treatment-associated response vector. Thus, the x and y axes differ only in the cellular context in which the virtual perturbation was applied. Points above the diagonal indicate targets whose predicted perturbation aligns more strongly with the treatment-associated direction in rheumatoid-arthritis patient cells than in healthy donor CD4+ T cells; points below the diagonal indicate targets with stronger alignment in the healthy donor context. Dot color indicates pathway membership: orange, TCR-proximal; blue, JAK-STAT; gray, other.

## Discussion

ImmunoSTATE benchmarks empirical and virtual perturbations against patient-derived transcriptomic trajectories in rheumatoid arthritis, yielding two principal findings. First, unbiased genome-wide screening consistently places TCR-proximal signaling above the measurable components of the JAK-STAT pathway as a transcriptomic reversal axis, suggesting that the molecular ground truth of patient improvement may extend beyond the targets currently exploited by standard-of-care therapies. Second, comparing virtual perturbation scores across healthy donor and rheumatoid-arthritis cellular contexts reveals that most target priorities are broadly preserved, yet a subset of targets, most notably SOX4, shows selective enrichment in the disease context that would not be apparent from healthy-donor data alone.

The empirical CRISPRi screen independently identified T cell-receptor-proximal signaling as the strongest route for reversing the measured rheumatoid-arthritis transcriptional program in primary T cells. Genes such as *ZAP70, CD3E, CD3G, LCP2, LCK, ITK,* and *LAT* ranked near the top of the genome-wide screen, whereas the measurable components of the JAK-STAT pathway were collectively ranked much lower. This result is biologically plausible given the central role of chronic antigen-driven T cell activation in sustaining pathogenic helper-like synovial states. At the same time, this finding should not be over-interpreted as definitive evidence that deep T cell-receptor blockade is the best clinical strategy. Our screen measures transcriptomic reversal in an experimental perturbation context, not organism-level efficacy or safety, and the direct tofacitinib targets *JAK1* and *JAK3* were not observable in the primary T cell atlas because of essential-gene dropout. The appropriate conclusion is therefore that T cell-receptor-proximal signaling outperformed the measurable components of the JAK-STAT pathway in this empirical benchmark.

Within this framework, virtual modeling is best viewed not as a replacement for empirical screening but as a contextual and hypothesis-generating layer. Applying perturbations directly to patient-derived pretreatment cells makes it possible to ask whether predicted shifts point in the same direction as the transcriptomic trajectory of clinical improvement. The strongest support in our study was confined to targets represented in the model training distribution. JAK1 and JAK3, which were absent from the training set, produced near-zero perturbation magnitudes and very broad uncertainty intervals and therefore should not be interpreted as meaningful virtual successes or failures. PRDM1, also absent from the training set, yielded a more stable negative estimate but remains out-of-distribution and should likewise be interpreted cautiously. This distinction is informative because it separates limitations of the model’s training distribution from the question of whether the model preserves target-specific signal for targets it has actually learned.

The context-comparison analysis further clarified what is gained by applying ImmunoSTATE to patient cells rather than relying only on healthy donor T-cell inference. Virtual target priorities in healthy donor CD4+ T cells and rheumatoid-arthritis CXCL13+ T cells were significantly concordant, indicating that ImmunoSTATE preserves a shared CD4+ T-cell prioritization structure across contexts. At the same time, the rheumatoid-arthritis patient-cell context reweighted specific targets in biologically interpretable ways. *SOX4* showed the clearest amplification in the disease context, consistent with prior evidence linking *SOX4* to CXCL13-producing helper T-cell programs under inflammatory conditions^30^. *IRF4* and *ITK* remained strong in both contexts, consistent with a context-stable T-cell activation and T-cell-receptor-proximal axis^29,31^. In contrast, *STAT4, RORC, TOX2*, and *BCL6* were attenuated in rheumatoid-arthritis patient cells relative to healthy donor CD4+ T cells, suggesting that conventional Th1, Th17, or follicular-helper-like programs are not uniformly prioritized in the CXCL13+ disease-cell state^39,41,42,43^. Thus, ImmunoSTATE adds a patient-cell-context layer that separates shared T-cell reversal biology from rheumatoid-arthritis-context-enriched target signals.

A key conceptual contribution of this work is that empirical and virtual perturbation analyses should not be treated as interchangeable readouts. The empirical screen provides an experimentally measured, genome-scale view of perturbation effects in primary human CD4+ T cells. The virtual analysis provides a patient-contextualized view of how selected targets are predicted to move rheumatoid-arthritis CXCL13+ T cells relative to the clinical improvement direction. The partial overlap between these two views is informative: shared targets may represent broadly conserved T-cell reversal biology, whereas targets amplified in the patient-cell context may reflect disease-state features not fully captured by healthy donor perturbation systems. In this sense, ImmunoSTATE is best understood as a framework for contextualizing empirical perturbation biology within patient-derived disease states, rather than as a replacement for experimental screening.

Several limitations and future directions should be considered. The patient-derived benchmark was built from nine patients treated with two therapeutic classes, so larger cohorts will be needed to separate shared treatment-associated directions from drug-specific or patient-specific response programs. The empirical perturbation arm used healthy donor primary CD4+ T cells, whereas the virtual arm was applied to rheumatoid-arthritis CXCL13+ synovial T cells; direct empirical perturbation data in patient-derived disease cells would provide the strongest validation. The STATE perturbation model was fine-tuned on a CD4+ T-cell perturbation atlas, so predictions for targets represented in the training distribution are more interpretable than predictions for targets absent from that training set. Future work should therefore combine patient-level pseudo bulk or mixed-effects modeling, therapy-specific benchmarks, direct perturbation assays in rheumatoid-arthritis T-cell states, and orthogonal functional readouts such as activation-marker and cytokine profiling. These studies will be particularly important for experimentally testing *SOX4*-associated CXCL13+ helper-cell biology and T-cell-receptor-proximal candidates such as *ITK* and *SYK*.

In summary, ImmunoSTATE provides a patient-anchored strategy for reading empirical and virtual perturbation data through the lens of clinical transcriptomic improvement. The framework identifies T-cell-receptor-proximal signaling as a strong empirical reversal axis, reveals interpretable virtual target signals in rheumatoid-arthritis patient cells, and shows that patient-cell context can reweight target priorities beyond what is apparent from healthy donor T-cell inference alone.

## Materials and Methods

### Clinical cohort and definition of a patient-derived treatment-associated transcriptomic reference

Single-cell RNA-sequencing (scRNA-seq) data from rheumatoid arthritis (RA) synovial fluid were obtained from the Gene Expression Omnibus (accession GSE296117)^9^. The cohort comprised nine patients with paired pre-treatment and post-treatment samples, including four treated with the JAK inhibitor tofacitinib^2^ and five with the TNF inhibitor adalimumab^3^.

Pathogenic CXCL13+ T cells^16^ were identified by first isolating CD4+ T cells on the basis of canonical lineage markers (*CD3D, CD3E*^44^*, CD4*^45^*, IL-7R*^46^*, TCF7*^47^, and *LEF1*^47^), followed by scoring each cell against a curated 14-gene pathogenicity module (CXCL13^16^, PDCD1^48^, MAF^35^, TOX^36^, TOX2^42^, BATF^49^, ICOS^50^, TIGIT^51^, IL21^52^, BCL6^43^, CXCR5^53^, SH2D1A^54^, CD200^55^, and SLAMF6^56^) using scanpy.tl.score_genes^57^. Cells exceeding the 90th percentile of the composite score were retained, yielding 1,307 pre-treatment and 3,600 post-treatment cells for downstream analysis. We refer to this population as CXCL13-associated pathogenic T cells because the selection module included CXCL13 together with additional peripheral-helper-like and dysfunction-associated markers, rather than relying on CXCL13 expression alone.

All cells were embedded into the 2,058-dimensional SE-600M^12^ latent space. The primary clinical reference used throughout the study was defined as the mean latent-space shift from the pre-treatment state to the post-treatment state across the analyzed CXCL13+ T cells. As patients receiving tofacitinib and adalimumab were analyzed jointly, this vector should be interpreted as a shared treatment-associated transcriptomic shift rather than a drug-specific response signature.

Differential expression between pre- and post-treatment cells was performed using the Wilcoxon rank-sum test in Scanpy v1.9^57^, with treatment-associated genes defined by a false discovery rate (FDR) below 0.05 and an absolute log_2_ fold-change exceeding 0.5, yielding 2,748 differentially expressed genes. Unless otherwise stated, all subsequent analyses were conducted at the single-cell level.

### Genome-wide perturbation screening and Reversal Gene Expression Score

Transcriptomic reversal potential was assessed genome-wide using the primary human CD4+ T cell CRISPR interference (CRISPRi) atlas generated by Zhu et al^7^. To ensure reliable interpretation of gene silencing effects, analysis was restricted to 9,085 genes for which CRISPRi achieved effective knockdown, defined as a knockdown effect size below -0.5 as reported by Zhu et al^7^.

For each perturbation *i*, reversal potential was quantified using the Reversal Gene Expression Score (RGES)^22^, which measures the degree to which silencing a given gene shifts the cellular transcriptome toward the post-treatment state. Let U denote genes upregulated in the pre-treatment state relative to the post-treatment state, and let D denote genes downregulated in the pre-treatment state relative to the post-treatment state, and let Δ*ig* denote the perturbation-induced expression change for gene *g* under perturbation *i*. RGES was then calculated as follows:

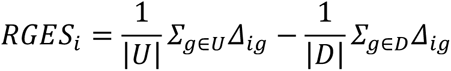

More negative values indicate that silencing gene *i* more strongly reverses the RA-associated transcriptional program toward the treated state. Perturbations were ranked by RGES, and pathway-level enrichment was assessed by testing whether genes belonging to a given signaling pathway ranked significantly higher than the genomic background using the Mann-Whitney U test^59^, with multiple testing correction applied using the Benjamini-Hochberg procedure^58^. To confirm that rankings were not specific to a single stimulation condition, RGES-based perturbation rankings derived from the 48-hour and 8-hour stimulation settings in the primary human CD4+ T cell CRISPRi atlas^7^ were compared using Spearman correlation.

Statistical significance of RGES was estimated using a one-sided permutation test in the reversal direction. For each perturbation, the observed RGES was compared with a null distribution generated by permuting the treatment-associated gene labels 10,000 times while preserving the sizes of the upregulated and downregulated gene sets. Empirical P values were calculated with add-one smoothing and adjusted across perturbations using the Benjamini-Hochberg procedure^58^. Because the screen was designed to prioritize perturbations that move cells toward the post-treatment reference, more negative RGES values were treated as more extreme for the primary ranking analysis.

### Leave-one-patient-out cross-validation

To assess the stability of the patient-derived reference and downstream target rankings, all analyses were repeated using nine-fold leave-one-patient-out (LOPO) cross-validation. In each fold, one patient was withheld and the gene expression signature representing the shared transcriptomic response to treatment was re-derived from the remaining eight patients using identical preprocessing and thresholds. The complete Gene-set Perturbation Scoring (GPS) pipeline, from RGES calculation to pathway-level ranking, was then repeated for each fold.

To confirm that findings were not driven by any individual patient, global ranking stability was quantified by Kendall’s rank correlation coefficient (τ) ^28^ between the ranking derived from each LOPO fold and the ranking derived from all nine patients combined. The relative ranking of TCR-proximal versus JAK-STAT signaling genes was reassessed within each fold using the Mann-Whitney U test^59^ to verify that pathway-level conclusions were preserved regardless of which patient was withheld. For a set of therapeutically relevant candidate genes, ranking consistency across the nine folds was further summarized using the coefficient of variation and the rank range across folds.

### Virtual perturbation modeling with STATE

Virtual perturbation predictions were generated using STATE^12^, an autoregressive transformer-based foundation model. SE-600M is a fixed encoder to embed single-cell transcriptomes into a 2,058-dimensional latent representation and was not fine-tuned in this study. Virtual perturbation predictions were generated with the ST-Tahoe perturbation transformer within the STATE framework. ST-Tahoe was fine-tuned on the primary human CD4+ T cell CRISPRi atlas^7^, which contains 12,107 perturbations across 2.05 million cells, and then applied to patient-derived pre-treatment CXCL13-associated synovial T cells to predict post-perturbation latent embeddings. Full details of the model architecture and training procedure are described in the original publication^12^.

Nineteen RA-relevant targets were selected *a priori* from TCR-proximal signaling, JAK-STAT signaling, and transcriptional regulators of pathogenic CXCL13+ T cell programs, on the basis of literature support, pathway relevance, and pharmacological tractability (**Supplementary Table S1**.). *In silico* perturbations were applied directly to the 1,307 patient-derived pre-treatment CXCL13+ T cells. For each target, a perturbation vector was computed as the mean shift in cell state between the model’s predicted post-knockdown representation and the unperturbed baseline, averaged across all 1,307 patient-derived pre-treatment cells. This vector captures the direction and magnitude of the predicted transcriptomic response to gene silencing.

Of the 19 targets, 16 were represented in the model training distribution and classified as trained targets. The remaining three (*JAK1*, *JAK3*, and *PRDM1*) were not present in the model’s training data. For these genes, the model did not use a learned target-specific representation, and their outputs were therefore interpreted cautiously rather than compared directly with trained targets.

### Evaluation of virtual perturbations in SE-600M latent space

The primary evaluation metric for virtual perturbation predictions was the cosine similarity between each target-specific perturbation vector and the patient-derived treatment-associated response vector in the 2,058-dimensional embedding space of SE-600M^12^, the single-cell foundation model that serves as the encoder within STATE and represents each cell’s gene expression profile as a vector of 2,058 numerical values. For a given target, the perturbation vector was defined as the mean difference between the predicted perturbed embedding and the corresponding unperturbed embedding across pretreatment patient T cells. Positive cosine similarity indicates that the predicted perturbation points in the same direction as the treatment-associated clinical shift.

Uncertainty was quantified by 2,000 bootstrap iterations in which perturbed and unperturbed cells were resampled with replacement and cosine similarity was recomputed at each iteration. Two-sided 95% confidence intervals were derived using the percentile method, and a target was considered to show significant alignment if the interval excluded zero. As this procedure resamples cells rather than patients, these intervals reflect cell-level uncertainty within the analyzed cohort and should not be interpreted as patient-level confidence intervals.

### Cross-context comparison between healthy donor and rheumatoid-arthritis cells

For the cross-context comparison, ImmunoSTATE scores were computed using the same patient-derived treatment-associated reference direction in both cellular contexts. This reference direction was derived from GSE296117^9^ rheumatoid-arthritis patient CXCL13+ T cells and was not re-estimated from the Zhu healthy-donor CD4+ T-cell atlas^7^. For each target represented in the model training distribution, virtual perturbation vectors were computed separately in healthy donor CD4+ T cells and in rheumatoid-arthritis patient CXCL13+ T cells. Each context-specific perturbation vector was then compared with the same patient-derived reference direction by cosine similarity. This design isolates the effect of cellular context on the predicted perturbation vector while holding the clinical reference direction fixed.

### Auxiliary validation analyses and negative controls

To assess whether the RA reversal signature captured T cell-specific biology rather than generic response to gene silencing, the same disease-associated signature was projected onto an orthogonal CRISPRi dataset generated in K562 chronic myeloid leukemia cells^23^, a cell line unrelated to the pathogenic T cell populations of interest. This negative-control dataset comprised 1,383 perturbations across 188,590 cells. Pseudobulk log_2_ fold-changes were computed per perturbation, RGES^22^ was recalculated, and overlap between the top 100 perturbations in the K562 and primary T cell analyses was quantified by direct intersection.

Biological concordance of the empirical prioritization framework was further examined using the Schmidt cytokine-conditioned perturbation benchmark^24^, a published primary-human-T cell CRISPR activation/interference resource spanning various cytokine and cytokine-pathway stimulation contexts. Directional concordance was evaluated by asking whether perturbation contexts expected to move the rheumatoid-arthritis program toward improvement ranked toward the favorable end of the benchmark, whereas opposite-direction contexts ranked toward the opposite end. For the auxiliary cytokine-conditioned benchmark, four rheumatoid-arthritis-relevant modules—pathogenicity, tissue destruction, inflammation, and B-cell help—were scored in each cytokine context, and an unweighted mean delta composite score was calculated. Eight clinically relevant contexts were prespecified for directional-concordance analysis. Pro-inflammatory cytokine blockade contexts were expected to rank toward the favorable end of the benchmark, whereas anti-inflammatory or opposite-direction control contexts were expected to rank toward the opposite end. Statistical support for the observed directional concordance was evaluated using five complementary tests: random sampling, empirical-base-rate testing, an exact binomial test, label permutation, and a rank-based test. These analyses were treated as supportive biological-concordance checks rather than primary inferential endpoints.

For pathway-level empirical comparisons, canonical T cell-receptor-proximal and JAK-STAT pathway members were curated from the literature and annotated for observability in the source primary-T cell CRISPRi atlas, including essential-gene-dropout events that precluded direct measurement of some loci. Full pathway gene lists and observability annotations are provided in **Supplementary Table S2**.

### Statistical analysis

Unless otherwise stated, all statistical tests were two-sided. The main exception was the RGES significance used for ranking in the empirical screen, which employed the one-sided permutation test described above because the objective was to identify perturbations that moved cells toward the post-treatment reference. Figure 2A presents a two-sided descriptive visualization of the same screen, but panels 2B-2D and the corresponding pathway-level conclusions are based on the one-sided reversal-oriented statistics. Multiple-testing correction was performed using the Benjamini-Hochberg procedure^58^ to control the false discovery rate. Bootstrap-derived confidence intervals are reported as percentile-based 95% intervals. Summary statistics are reported as mean ± standard deviation unless otherwise indicated. Because the primary treatment-associated reference and the main uncertainty estimates were derived from cell-level data within a small, paired cohort, statistical results should be interpreted as robustness analyses within the analyzed dataset rather than as definitive estimates of population-level patient effects.

## DATA AVAILABILITY

The patient scRNA-seq data analyzed in this study are available in Gene Expression Omnibus under accession GSE296117^9^. The primary human CD4+ T cell CRISPRi atlas is from Zhu *et al.*^7^ The orthogonal K562 Perturb-seq dataset is from Replogle *et al*.^23^ The STATE model is described by Adduri *et al*.^12^

## ACKNOWLEDGEMENTS

This work was supported by KAKENHI grants from the Japan Society for the Promotion of Science (JSPS) to H.S. (23K28184, 24H01755 and 25H01571), JST FOREST Program to H.S. (JPMJFR242Q), as well as the Canon Foundation and Nakatani Foundation. Figures were created in part with BioRender.com. We thank K. Tanaka for assistance with manuscript preparation.

## AUTHOR CONTRIBUTIONS

K.N. initially conceived of and designed the project. H.S. supervised the study. K.N. performed all formal analyses. K.N. and H.S. jointly wrote the manuscript. All authors have read and approved the final manuscript.

