## Supplementary Tables S1, S2, Supplementary Figures S1, S2, S3 for "Integrating virtual cell modeling with patient-derived transcriptomics to uncover target signals in rheumatoid arthritis"

**Supplementary Table S1 | Nineteen rheumatoid-arthritis-relevant targets used for virtual perturbation modeling**

| **Target** | **Class** | **Selection rationale and training status** |
|---|---|---|
| *TOX* | Transcription factor | Candidate regulator of pathogenic helper-like and dysfunction-associated T cell states; trained target |
| *SOX4* | Transcription factor | Candidate regulator of CXCL13-positive helper-like T cell identity; trained target |
| *MAF* | Transcription factor | B cell-help and peripheral-helper/T-follicular-helper program; trained target |
| *BATF* | Transcription factor | Pathogenic helper-like transcriptional program; trained target |
| *BCL6* | Transcription factor | Canonical helper/T-follicular-helper regulator; trained target |
| *PRDM1* | Transcription factor | Differentiation regulator in helper-like T cell states; absent from the model training set; interpreted cautiously |
| *EOMES* | Transcription factor | Effector or dysfunction-associated transcriptional regulator; trained target |
| *TBX21* | Transcription factor | Inflammatory T cell lineage regulator; trained target |
| *RORC* | Transcription factor | Inflammatory T cell lineage regulator; trained target |
| *IRF4* | Transcription factor | Candidate regulator of pathogenic helper-like synovial T cell programs; trained target |
| *TOX2* | Transcription factor | TOX-family regulator linked to dysfunctional T cell programs; trained target |

| | | |
|---|---|---|
| *JAK1* | Kinase | Clinically targeted Janus kinase in rheumatoid arthritis; absent from the model training set; interpreted cautiously |
| *JAK3* | Kinase | Clinically targeted Janus kinase in rheumatoid arthritis; absent from the model training set; interpreted cautiously |
| *BTK* | Kinase | Druggable kinase with emerging autoimmune relevance; trained target |
| *SYK* | Kinase | Druggable kinase with established pathway relevance in inflammatory signaling; trained target |
| *ITK* | Kinase | T cell-receptor-proximal kinase with pathway relevance in pathogenic T cells; trained target |
| *STAT3* | Signaling node | JAK-pathway downstream mediator and measurable comparator target; trained target |
| *STAT4* | Signaling node | JAK-pathway downstream mediator and measurable comparator target; trained target |
| *NFKB1* | Signaling node | Canonical inflammatory signaling hub relevant to rheumatoid arthritis; trained target |

**Supplementary Table S2 | Canonical T cell-receptor-proximal and JAK-STAT pathway members used for the empirical pathway comparison**

Pathway membership was curated from rheumatoid-arthritis and T cell-signaling literature; “Observable” indicates whether a gene was directly measurable in the primary human CD4+ T cell CRISPRi atlas used for GPS.

A. T cell-receptor-proximal signaling genes (n = 9)

| Gene | Functional tier | Observable in primary CD4+ T cell CRISPRi atlas | Included in empirical pathway comparison | Notes |
|---|---|---|---|---|
| *ZAP70* | Kinase | Yes | Yes | Observable; included in empirical comparison |
| *CD3E* | Receptor complex | Yes | Yes | Observable; included in empirical comparison |
| *CD3G* | Receptor complex | Yes | Yes | Observable; included in empirical comparison |
| *CD3D* | Receptor complex | Yes | Yes | Observable; included in empirical comparison |
| *LCP2* | Adaptor / scaffold | Yes | Yes | Observable; included in empirical comparison |
| *LAT* | Adaptor / scaffold | Yes | Yes | Observable; included in empirical comparison |
| *LCK* | Kinase | Yes | Yes | Observable; included in empirical comparison |
| *ITK* | Kinase | Yes | Yes | Observable; included in empirical comparison |
| *FYN* | Kinase | Yes | Yes | Observable; included in empirical comparison |

B. JAK-STAT pathway genes (n = 20)

| Gene | Functional tier | Observable in primary CD4+ T cell CRISPRi atlas | Included in empirical pathway comparison | Notes |
|---|---|---|---|---|
| JAK1 | Kinase | No | No | Not observable in the primary CD4+ T cell CRISPRi atlas; excluded because of essential-gene dropout |
| JAK2 | Kinase | Yes | Yes | Observable; included in empirical comparison |
| *JAK3* | Kinase | No | No | Not observable in the primary CD4+ T cell CRISPRi atlas; excluded because of essential-gene dropout |
| *TYK2* | Kinase | Yes | Yes | Observable; included in empirical comparison |
| *STAT1* | STAT transcription factor | Yes | Yes | Observable; included in empirical comparison |
| *STAT2* | STAT transcription factor | Yes | Yes | Observable; included in empirical comparison |
| *STAT3* | STAT transcription factor | Yes | Yes | Observable; included in empirical comparison. Only measurable JAK-STAT member in the top 100 |
| *STAT4* | STAT transcription factor | Yes | Yes | Observable; included in empirical comparison |

| Gene | Functional tier | Observable in primary CD4+ T cell CRISPRi atlas | Included in empirical pathway comparison | Notes |
|---|---|---|---|---|
| *STAT5A* | STAT transcription factor | Yes | Yes | Observable; included in empirical comparison |
| *STAT5B* | STAT transcription factor | Yes | Yes | Observable; included in empirical comparison |
| *STAT6* | STAT transcription factor | Yes | Yes | Observable; included in empirical comparison |
| *IRF1* | Interferon-linked transcription factor | Yes | Yes | Observable; included in empirical comparison |
| *IRF9* | Interferon-linked transcription factor | Yes | Yes | Observable; included in empirical comparison |
| *SOCS1* | Negative regulator | Yes | Yes | Observable; included in empirical comparison |
| *SOCS3* | Negative regulator | Yes | Yes | Observable; included in empirical comparison |
| *CISH* | Negative regulator | Yes | Yes | Observable; included in empirical comparison |
| *PIAS1* | Negative regulator | Yes | Yes | Observable; included in empirical comparison |
| *PTPN1* | Phosphatase / regulator | Yes | Yes | Observable; included in empirical comparison |

| Gene | Functional tier | Observable in primary CD4+ T cell CRISPRi atlas | Included in empirical pathway comparison | Notes |
|---|---|---|---|---|
| *PTPN2* | Phosphatase / regulator | Yes | Yes | Observable; included in empirical comparison |
| *PTPN11* | Phosphatase / regulator | Yes | Yes | Observable; included in empirical comparison |

**Notes.** All nine T cell-receptor-proximal genes were observable in the source primary human CD4$^{+}$ T cell CRISPRi atlas and were included in the pathway-level empirical comparison. Among the 20 canonical JAK-STAT pathway members, 18 were observable; *JAK1* and *JAK3* were not directly measurable and were excluded because of essential-gene-dropout in the empirical atlas. In the empirical screen, *STAT3* was the only measurable JAK-STAT member entering the top 100, whereas the remaining 17 measurable JAK-STAT members ranked 1,457th or lower.

### Supplementary figure legends

**Supplementary Figure S1 | One-sided reversal-oriented view of the empirical screen**

One-sided reversal-oriented view of the same RGES distribution summarized descriptively in Figure 2A. Here the y-axis shows the one-sided false-discovery rate (FDR) for movement toward the post-treatment reference, so that only perturbations shifting cells toward the treated state are scored favorably. By construction, perturbations that shift cells away from the post-treatment state lie near $\mathrm{FDR} \approx 1$ even when their absolute effect size is large. The strongest therapeutic cluster on the negative-RGES side is enriched for TCR -proximal genes (orange), with JAK-STAT genes (blue) and all other perturbations (gray) shown for reference. Because the empirical *P* value cannot be smaller than$1/10{,}000$ given the permutation depth, the most significant hits form a tied ceiling after Benjamini-Hochberg correction. This one-sided screen was used for pathway ranking and therapeutic prioritization in Figure 2B-D.

**Supplementary Figure S2 | Orthogonal K562 negative-control analyses**

**A, Minimal overlap between K562 and T cell top-ranked perturbations.** The number of genes shared between the top 100 RGES-ranked perturbations in K562 cells and in primary CD4+ T cells is shown. Only four genes were common to both lists, confirming the strong cell-type specificity of the RA reversal signature.

**B, Cell-type specificity of the RA reversal signature.** RGES-ranked perturbations from the K562 chronic myeloid leukemia CRISPRi dataset scored against the RA reversal signature. In contrast to the TCR-proximal signaling genes that dominate the top rankings in primary $CD4^+$ T cells, the leading K562 hits were enriched for RNA-processing and splicing factors (for example, *SMG5, AQR, HNRNPC*, and *SLU7*) and Integrator-complex members (*INTS2, INTS8, and INTS5*). Gene-name colors indicate functional class: red, RNA processing and splicing; blue, Integrator complex; gray, other housekeeping genes.

**Supplementary Figure S3 | Orthogonal biological-concordance checks using the Schmidt cytokine-conditioned perturbation benchmark.**

**A,** Directional concordance across prespecified cytokine contexts. The Schmidt primary human T cell CRISPR activation and interference resource was reanalyzed by scoring four RA-relevant gene modules—pathogenicity, tissue destruction, inflammation, and B cell help—and averaging their delta scores into a composite cytokine-response benchmark. All 91 cytokine or cytokine-pathway contexts in the resource were ranked by this composite score, where lower rank indicates stronger concordance with the patient-derived RA improvement direction. Eight clinically relevant contexts were highlighted, including blockade of IL-1β, GM-CSF, IL-17A, IL-6, and TNF-α, together with IL-10, IL-2, and TRAIL stimulation controls. Orange bars indicate pro-inflammatory blockade contexts that followed the expected directional pattern, teal bars indicate anti-inflammatory or opposite-direction contexts that ranked toward the unfavorable end as expected, and the gray bar indicates the single pro-inflammatory discordant context. IL-10, IL-2, and TRAIL contexts ranked at the unfavorable end of the distribution as expected, given that these represent anti-inflammatory or non-therapeutic stimulation conditions rather than as favorable therapeutic perturbations.

**B,** Null-model support for the observed directional concordance. Statistical evaluation of the 7-of-8 directional concordance observed in panel A. Bars show $-\log10(P)$ values from five complementary tests evaluating whether the observed 7-of-8 directional concordance exceeds chance expectation. Tests comprised random sampling of eight contexts from the 91-condition universe, an empirical-base-rate test using the observed favorable-direction frequency as the null, an exact binomial test with $p = 0.5$, permutation of expected-direction labels across the observed delta scores, and a rank-based test comparing the pro-inflammatory blockade contexts against random draws. The dashed horizontal line marks the nominal significance threshold at $\alpha = 0.05$. These analyses therefore provide supportive, but not uniformly definitive evidence for biological concordance of the empirical prioritization framework.

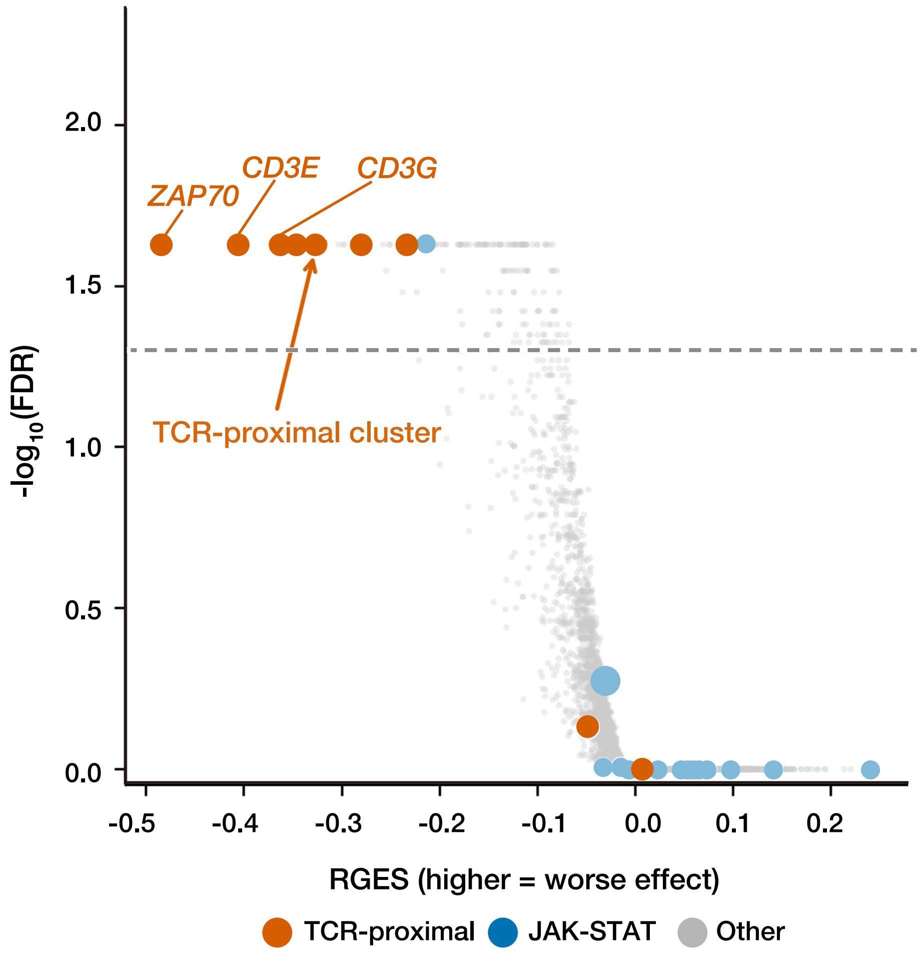


**Nakanishi *et al.* Supplementary Figure S1**

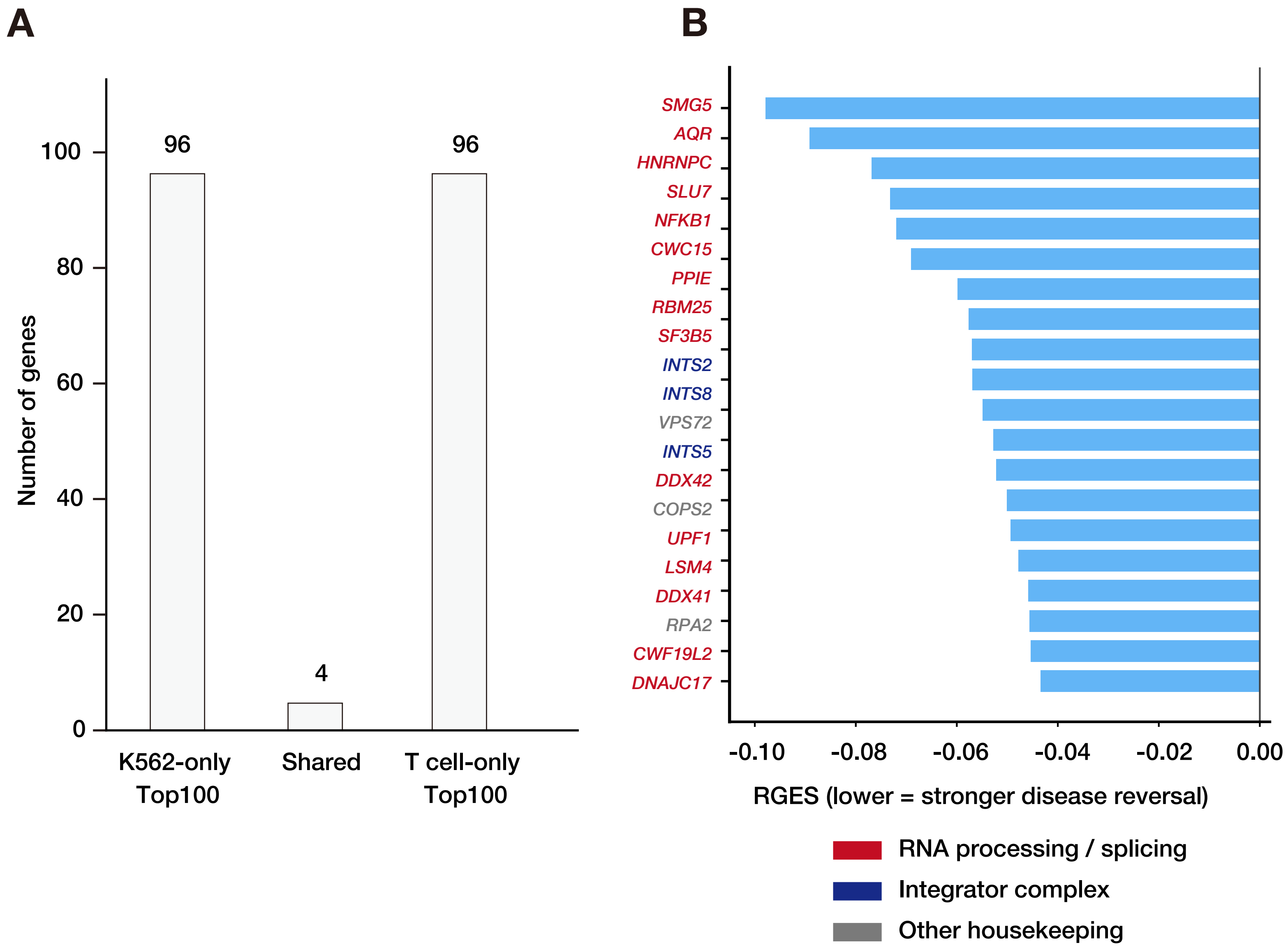


**Nakanishi *et al.* Supplementary Figure S2**

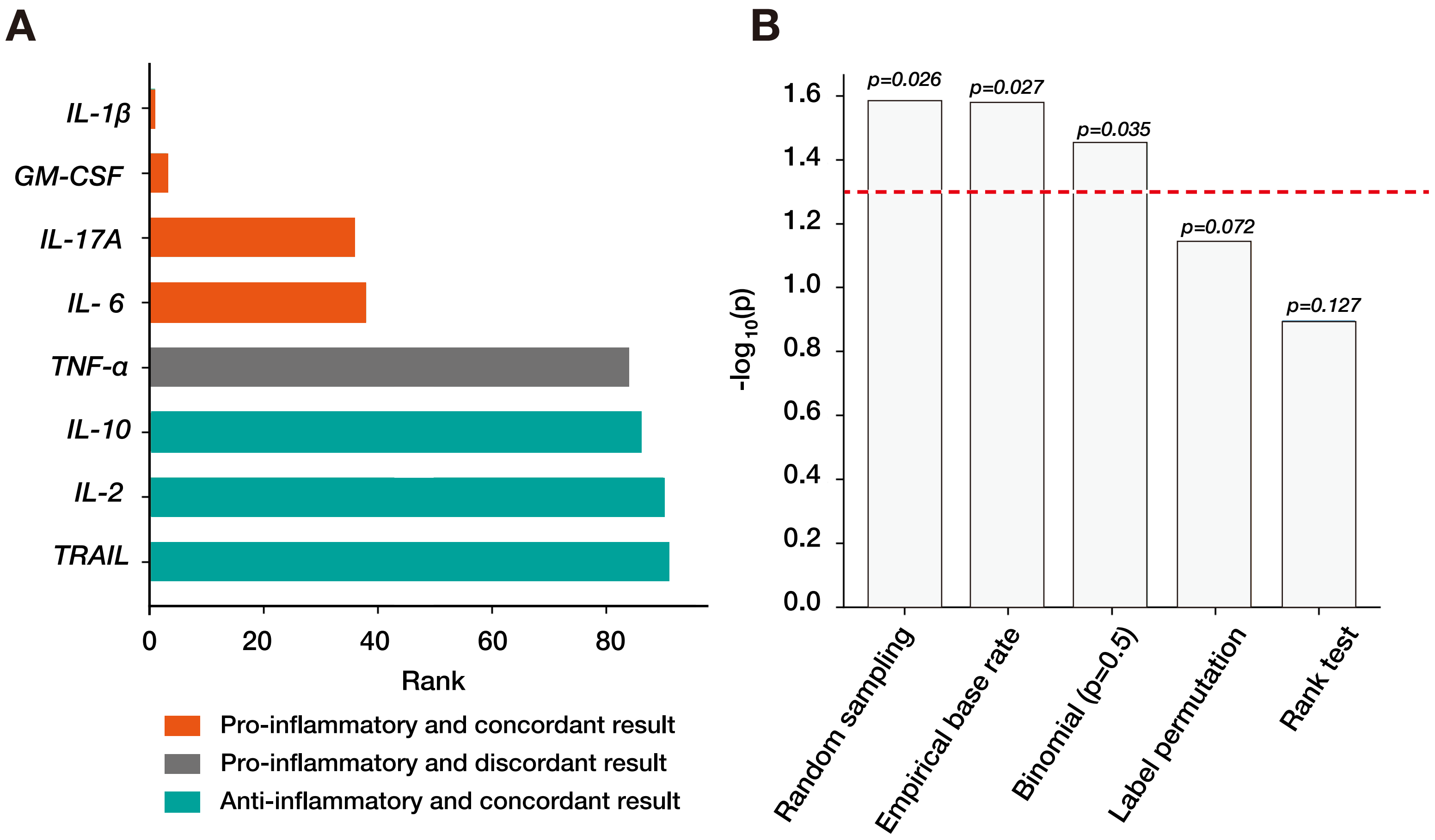


**Nakanishi *et al.* Supplementary Figure S3**
